# Distinct Hippocampal Cellular Pathologies Influence Cognition Across Diagnostic Categories and Distinguish Schizophrenia from Affective Psychosis

**DOI:** 10.64898/2026.04.27.720978

**Authors:** Eugene Ruby, Oded Gonen, Julie Walsh-Messinger, Eyal Lotan, Assaf Tal, Henry Rusinek, Jose C. Clemente, Jessica Robinson-Papp, Ezequiel Lafont, Katherine H. Karlsgodt, Dolores Malaspina

**Author notes:** **Corresponding Author:** Dolores Malaspina, Department of Psychiatry at Icahn School of Medicine at Mount Sinai, 1 Gustave L. Levy Place, Box 1230, New York, NY 10016.

## Abstract

Cognitive deficits are core features of psychosis and strong predictors of functional outcome, but are largely refractory to pharmacological treatments. It remains unresolved whether these deficits share cellular mechanisms across the psychosis spectrum (as in transdiagnostic models) or differ across DSM-5 diagnostic boundaries, particularly those separating schizophrenia from affective psychoses. We tested this by examining associations between MATRICS Cognitive Battery scores and whole-hippocampal proton magnetic resonance spectroscopic imaging (^1^H-MRSI) neurometabolite biomarkers of cellular pathologies in 33 participants with psychosis (20 schizophrenia, 13 affective psychosis), 22 with non-psychotic affective disorders, and 25 healthy controls. Metabolite biomarkers included N-acetylaspartate (NAA) for neuronal integrity; myo-inositol (mI) for glia; choline (Cho) for membrane/myelin turnover; and glutamate+glutamine (Glx) for glutamatergic neurotransmission. Metabolite concentrations did not differ between groups but showed group-specific associations with cognition. In healthy controls, lower NAA was associated with poorer overall cognition, working memory, and reasoning (r_s_=.41, .42, and .43, respectively; p’s<.05). Among non-psychotic affective cases, lower mI and higher Cho were associated with worse verbal learning (r_s_=.50 and −.45, respectively; p’s<.05). In psychosis, lower Cho predicted poorer visual learning (r_s_=.51; p=.002). The clearest between-group divergence involved social cognition, the cognitive domain most predictive of functioning in psychosis: worse performance was associated with lower Glx in schizophrenia (r_s_=.67; p=.033) but higher Cho and NAA in affective psychosis (r_s_=−.58 and −.58, respectively; p’s<.05). Each metabolite-social cognition association was opposite in direction and differed significantly between these groups (p’s<.05), contrary to what transdiagnostic models would predict. These findings suggest distinct hippocampal cellular substrates of social cognitive impairments in schizophrenia and affective psychosis, supporting diagnosis-specific treatment targets and warranting replication in larger samples.

## INTRODUCTION

Approximately 1% of the U.S. population has a schizophrenia-related psychotic disorder, most of whom do not experience a full remission. ^1^ Cognitive deficits are consistently observed in schizophrenia-related psychosis, ^2, 3^ including first-episode psychosis, ^4^ and thus impaired cognition is considered a core feature of the disease. While currently available antipsychotic medications are effective for reducing positive symptoms, no study has produced robust, clinically meaningful efficacy data for any agent to remediate cognitive impairments, ^5^ which predict worse functional outcomes and long-term disability. ^3^

A roadblock to identifying pharmacological treatments for cognition could be the common practice of grouping schizophrenia and affective psychosis patients together (e.g., in transdiagnostic research), as schizophrenia and other psychoses may have distinct cellular abnormalities for cognitive deficits. The question of whether the neuropathology of psychosis is similar in schizophrenia and affective psychosis harkens back to 1840, when Albert Zeller first proposed a unitary theory of psychosis that hypothesized a single cross-disorder pathology. ^6^ This remained the prevailing theory until nearly 50 years later when Emil Kraepelin proposed a ‘dichotomy’ between dementia praecox (schizophrenia) and manic-depressive insanity (bipolar disorder). In 2009, the NIMH built upon Zeller’s unitary approach in constructing the Research Domain Criteria (RDoC) framework. It proposes a transdiagnostic hypothesis of common underpinnings for each domain of psychopathology (including each cognitive domain), ranging from cellular and molecular abnormalities to circuitry dysfunction and varying only in severity across diagnostic classifications. ^7^ However, if the mechanisms underlying cognitive deficits are distinct for schizophrenia and affective psychosis, samples that combine both types of psychosis could confound study results.

To examine this possibility, we measured levels of metabolite biomarkers of cellular pathology using three-dimensional whole-hippocampal proton MR spectroscopic imaging (^1^H-MRSI) and examined their associations with MATRICS Cognitive Consensus Battery (MCCB) global and domain scores across diagnostic groups. The hippocampal metabolites used as biomarkers were N-acetylaspartate (NAA) for neuronal integrity; creatine (Cr) for cellular energy metabolism; choline (Cho) for membrane/myelin turnover; myo-inositol (mI) for glia; and glutamate plus glutamine (Glx) for glutamatergic neurotransmission. ^8^ The hippocampus was chosen because of its central role in cognition and its implications for schizophrenia. We first analyzed the associations in separate groups of persons with any psychoses, non-psychotic affective disorders, and healthy controls, and then repeated the analyses after separating the psychosis group into schizophrenia and affective psychosis subgroups. This allowed us to test whether there was a common pathology associated with cognitive deficits across diagnoses, including healthy persons not meeting criteria for any disorder. We hypothesized that associations between metabolite biomarkers and cognition would vary by clinical group in a domain-specific manner, including for schizophrenia versus affective psychosis.

## METHODS

### Human Subjects

Patients with psychosis (with and without affective symptoms) and non-psychosis affective disorders (NP-affective) were recruited from treatment settings at Mount Sinai Hospital for an NIMH-funded study investigating the potential role of hippocampal inflammation in the pathophysiology of psychosis. All patients were on stable medications for at least one month. Healthy controls, recruited from the local community via flyers and web postings, had no history of psychosis, did not meet criteria for any DSM-5 diagnosis in the preceding two years, had no family history of psychosis, and were not taking psychiatric medications. Exclusion criteria for all participants included severe head trauma, premorbid intellectual disability, suicide risk, seizures (other than medication-related), and MRI contraindications. The study was approved by the local Institutional Review Board (IRB) and all participants provided written informed consent.

### Clinical Procedures

The Diagnostic Interview for Genetic Studies (DIGS) ^9^ was administered by trained clinicians with a master’s degree or higher to determine DSM-5 diagnoses and to classify patients as having psychosis or a non-psychotic affective disorder. The psychosis group included participants with schizophrenia and those with affective psychosis(schizoaffective and bipolar I disorder with psychosis). Cognition was assessed with the MATRICS test battery, ^10, 11^ which yields age and sex-normed t-scores for the domains of working memory, attention/vigilance, verbal learning, visual learning, reasoning/problem solving, processing speed, and social cognition. These seven domain scores are also combined to produce an age and sex-normed overall cognition composite score.

### MRI and Hippocampal Spectroscopic Imaging – Acquisition

Imaging was done in a 3 T whole-body scanner (Skyra, Siemens AG, Erlangen, Germany) using a transmit-receive head-coil (TEM3000, MR Instruments, Minneapolis, MN). Subjects were placed head-first supine into the magnet, followed by sagittal *T_1_*-weighted magnetization-prepared rapid gradient-echo (MP-RAGE) MRI: *TE*/*TR*/*TI*=3.5/2150/1000 ms, 7° flip-angle, 160 slices 1 mm thick, 256×224 matrix, 256×256 mm^2^ field of view. These were reformatted into 192 axial, sagittal and coronal slices angled along the hippocampal long axis at 1 mm^3^ isotropic resolution, for ^1^H-MRSI volume-of-interest (VOI) placement and tissue segmentation, as shown in Fig. 1a,b. ^12^

**Figure 1.**
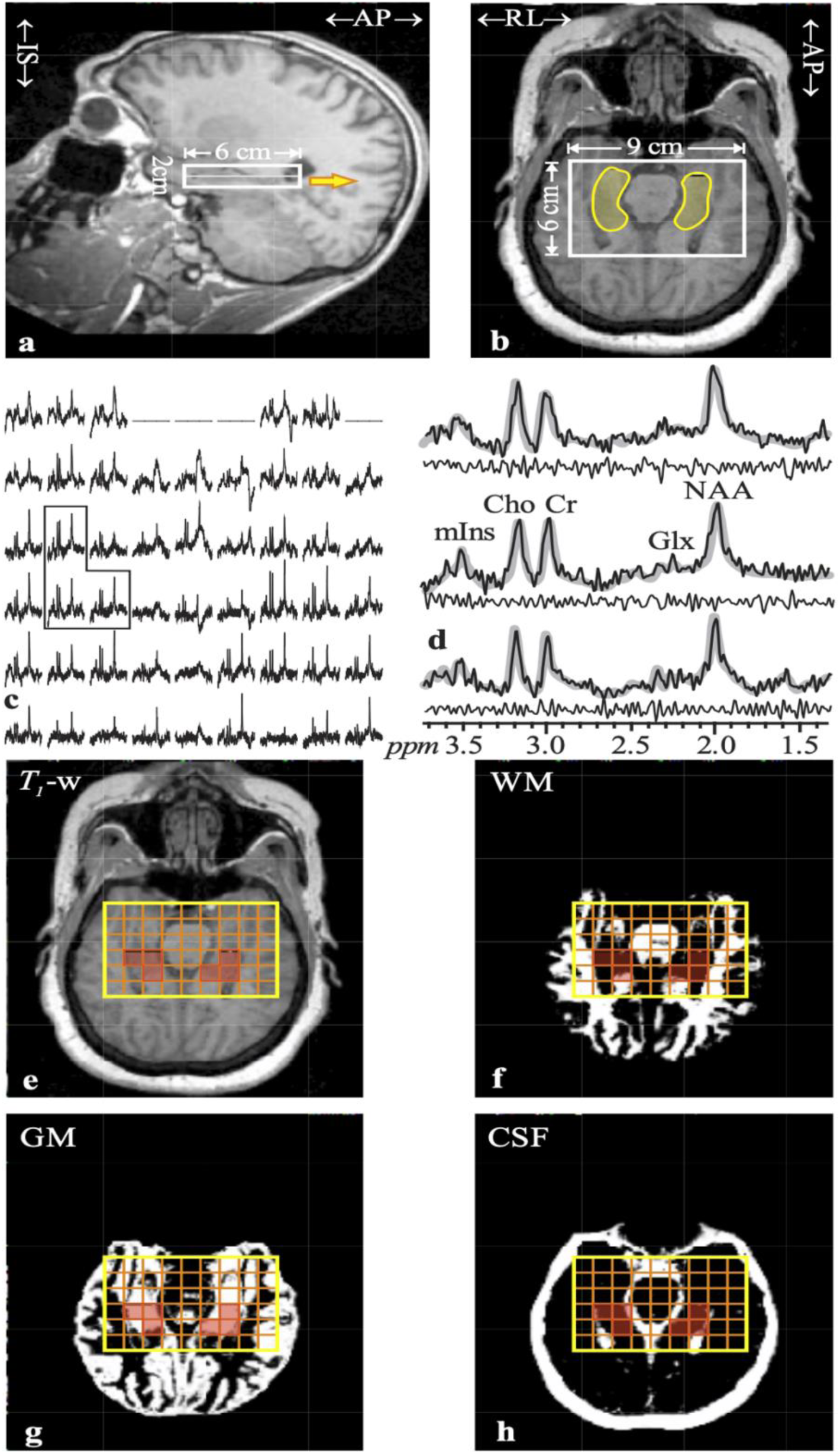
Imaging and ^1^H-MRSI Position, Size and Analysis. **Top:**Sagittal (**a**) and axial (**b**) *T_1_*-weighted MRI of a 22 year-old female with schizophrenia, superimposed with the 9×6×2 cm^3 1^H-MRSI VOI (thick white frames) and outline of hippocampus based on Frisoni guidelines (transparent yellow zones on **b**). Yellow arrow on **a** indicates the slice-level of **b**. **Center:** left - **c**: Real part of the 9×6 (LR×AP) axial ^1^H spectra matrix from the VOI slice on **b**. Spectra within the right hippocampus on **a** are marked by the dashed frame. Right – **d**: The spectra in that frame, expanded for detail (black lines) superimposed with the spectral fit (gray, see Fig. 2). Note the good signal-to-noise ratio; spectral resolution (8.1±3.0 Hz linewidth) from the (1 cm^3^) voxels; and fit fidelity, reflected by the “noise” (experimental – fit) residual below each spectrum. **Bottom: e – h**: Axial MRI (**e**), and its WM (**f**), GM (**g**), and CSF (**h**) probability maps, obtained with SPM12, superimposed with the ^1^H-MRSI grid (orange matrix) and the bilateral hippocampus outline (transparent red). These are used by our software to identify hippocampal voxels and extract their GM spectroscopic content (corrected for CSF and WM partial volumes), as described in the Methods section.

A 6 cm anterior-posterior (AP) ×9 cm left-right (LR) ×2 cm inferior-superior (IS)=108 cm^3 1^H-MRSI VOI was then image-guided over the bilateral hippocampus, as shown in Fig. 1a,b. The VOI was excited with point resolved spectroscopy (PRESS: *TE*/*TR*= 120/1500 ms). This intermediate *TE* was a compromise to *(i)* attenuate unwanted signals from short *T_2_* species (lipids and macromolecules) that were either within the VOI or not sufficiently excluded by the selective PRESS pulses; *(ii)* incur an acceptable *T_2_*signal loss of 25-30%; *(iii)* allow the *J*-coupled multiplets of Glx and mI to partially refocus, as shown in Fig. 2. The VOI was encoded into two axial 1 cm thick slices, and 12×12 gradient-phase-encoded in their planes, as shown in Fig. 1b, ^13, 14^ to form (1.0 cm)^3^ voxels. The ^1^H-MRSI signals were acquired for 256 ms at ±1 kHz bandwidth. At two averages the ^1^H-MRSI was ∼15 minutes and the entire protocol took under 40 minutes. ^12^

**Figure 2.**
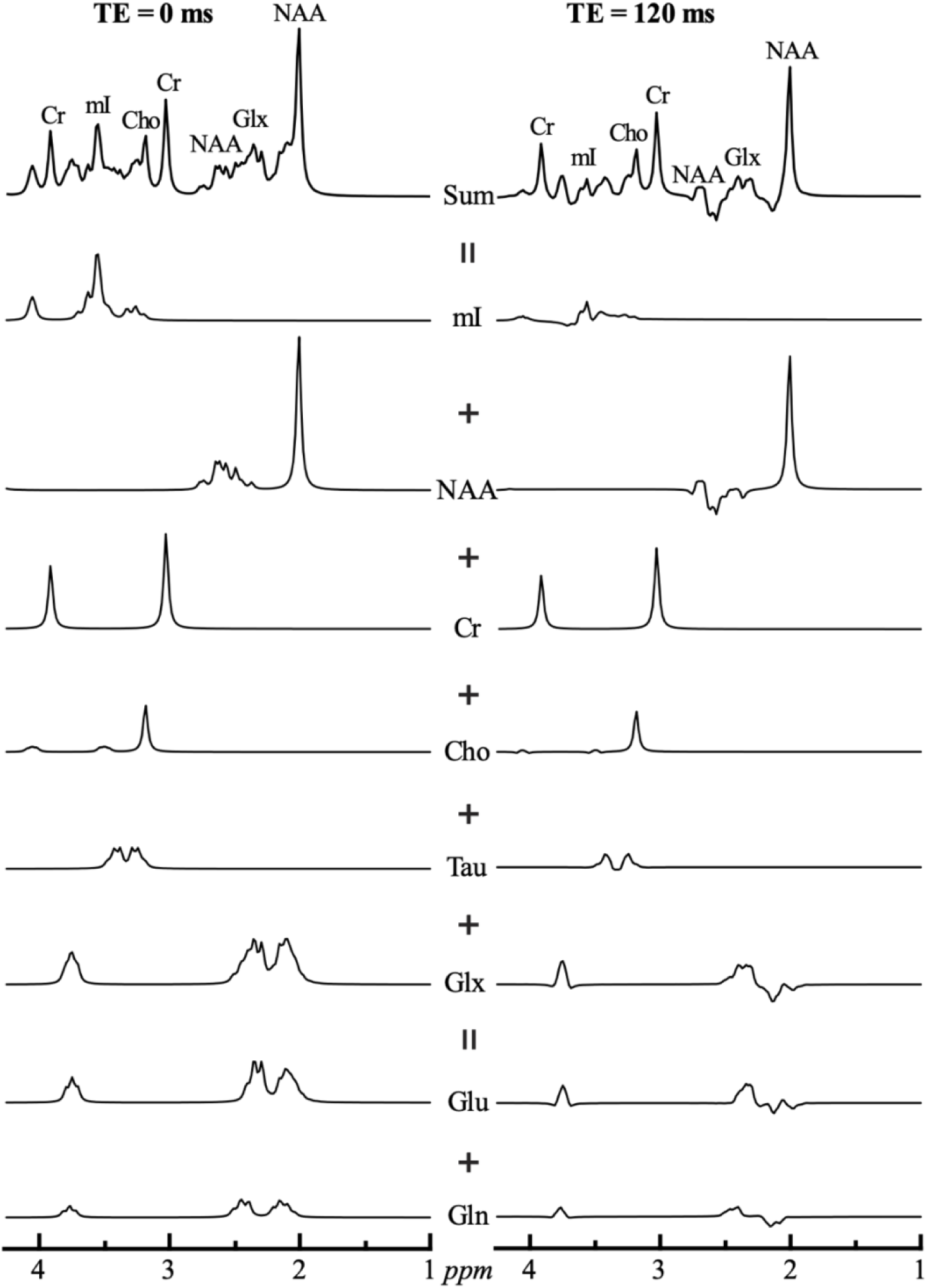
^1^H-MRSI Spectral Fitting Model Functions. Note. Simulated ^1^H-MRS spectra at *TE*=0 (left) and TE=120 ms (right), assuming metabolites’ *T_2_*=¥. These spectra were synthesized by summing the different simulated basis functions of the individual metabolites (**below**), generated at *TE=*0 ms (left) and *TE=*120 ms (right), obtained by full density matrix-based spectral simulation for each metabolite in the GAVA-Simulation application using the sequence’s actual complex RF pulse waveforms and timings.^45^ The waveforms are weighted by their reported *in vivo* concentrations expected in the central nervous system.^46^ NAA=10 mM, Cho=1 mM, Cr=6 mM, mI=6 mM, Glx=11.5 mM (Glu=10 mM + Gln=1.5 mM), Taurine=4 mM. Note: *(i)* the *J*-coupled multiplets (Glx and mI) partially refocus sufficiently to be distinct, even at *TE=*120 ms; *(ii)* the concordance of the full simulated *TE=*120 ms spectrum and the experimental spectra in Fig. 1d.

### MRI and Hippocampal Spectroscopic Imaging – metabolic quantification

^1^H-MRSI data were post-processed with in-house software (IDL 8.7.3, L3Harris Geospatial, Broomfield, CO). Residual water was removed from the signals in the time domain, ^15^ and the data were corrected for magnetic-field drift, voxel-shifted to align the localization grid with the NAA VOI, and zero-filled from 12×12 to 16×16 voxels in the slices’ planes and from 512 to 2048 in the time domain. The data were then Fourier transformed in the temporal, AP, and LR directions and Hadamard reconstructed along the IS. Each spectrum was automatically corrected for frequency and zero-order phase shifts, as shown in Fig. 1c,d.

Relative levels of each voxel’s NAA, Glx, Cr, Cho and mI were estimated from their spectral peak areas using the SiTToolsFit spectral modeling package^16^ with model functions for Glu+Gln (=Glx), Cho, Cr, mI, NAA, and taurine (Fig. 2), as shown in Fig. 1d. The relative levels were scaled into absolute levels against a 2 L reference sphere of known NAA, Cr, Cho and mI concentrations in water. ^17^

For tissue segmentation and ^1^H-MRSI quantification, bilateral (left + right) hippocampal masks were traced by a trained neuroradiologist on the sagittal images, according to the Harmonized Protocol of the European Alzheimer’s Disease Consortium, ^18, 19^ (see Fig. 1b). The axial MRI images were segmented into cerebrospinal fluid (CSF), gray matter (GM), and white matter (WM) masks with SPM12 [University College London, UK], ^20^ as shown in Fig. 1f-h. In-house software (MATLAB 18, MathWorks, Framingham, MA) retained for analyses only voxels meeting the following criteria: *(a)* at least 30% of their volume inside the hippocampal mask; *(b)*‹30% CSF; *(c)* Cramer-Rao lower bounds ‹20% for any metabolite; and *(d)* 4 Hz‹linewidths‹13 Hz (both *(c)* and *(d)* are more stringent than the MRS consensus criteria described by Kreis el., 2004, and Wilson *et al., 2019*), ^21, 22^ as shown in Fig. 1e-h. The metabolites’ average GM concentrations in the retained (*N*›2) hippocampi voxels were estimated by our software with linear regression, as described by Tal *et al*., 2012. ^23^

### Statistical Analyses

IBM SPSS Statistics (version 31) was used for statistical analyses. One-way ANOVA, Kruskal-Wallis test, or Welch’s ANOVA compared the groups on age, metabolite concentrations, and cognitive measures. Kruskal-Wallis test or Welch’s ANOVA were used when conventional ANOVA assumptions were violated, the latter specifically in cases of heterogeneity of variance given the unequal group sample sizes. ^24^ Mann-Whitney U and Kruskal-Wallis tests compared the psychiatric groups based on illness duration. Group differences in sex makeup were assessed with Chi-square test. One-way ANCOVA compared metabolite levels across the two psychiatric groups while controlling for age and sex, and MATRICS scores were age- and sex-corrected using the MATRICS scoring program. ^11^ Levene’s test was used to assess differences in the variance of each metabolite between groups. For each ANOVA/ANCOVA or equivalent non-parametric test, *post-hoc* comparisons for each group combination were performed with correction for multiple comparisons using Bonferroni correction to maintain α=.05. Within each group, Spearman correlations tested relationships between metabolite levels and cognition. Fisher’s r-to-z transformation was used to normalize each correlation coefficient to allow for comparisons of correlation coefficients between groups. We also examined the potential effects of age and illness duration on the metabolite-cognition associations by assessing correlations of cognitive scores and metabolite levels with age and illness duration in the overall psychosis group and the schizophrenia and affective psychosis subgroups.

## RESULTS

The 33 participants with psychosis, 22 with non-psychotic affective disorders, and 25 healthy controls did not significantly differ in age (p=0.575) or sex (p=0.138), and the clinical groups had similar illness durations (p=0.409; see Table 1). Similarly, after stratification of the psychosis group into subgroups, schizophrenia (n=20), affective psychosis (n=13), non-psychotic affective disorders, and healthy controls did not significantly differ in age (p=.220) or sex (p=.138), and the clinical groups had similar illness durations (p=.470). Comparisons between just schizophrenia and affective psychosis are also shown in Supplementary Table 1.

**Table 1.** Demographic, Cognitive, and Imaging Data Stratified by Participant Group.

|  | **Psychosis<br>(n = 33)** | **NP-affective<br>(n = 22)** | **Healthy<br>Controls<br>(n = 25)** | **Test Statistic<sup>a</sup>** | **p** |
|---|---|---|---|---|---|
| **Demographics** |  |  |  |  |  |
| Age, *M (SD)* | 35.79 (10.91) | 36.27 (11.82) | 32.44 (8.41) | H(2) = 1.11 | p = .575 |
| Disease duration (years), *M (SD)* | 18.48 (12.61) | 15.55 (10.97) | n/a | U = 315.00 | p = .409 |
| Sex *(Percent male:female)* | 51.5 : 48.5 | 27.3 : 72.7 | 32.0 : 68.0 | χ<sup>2</sup>(2) = 3.97 | p = .138 |
| **MATRICS Cognitive Battery** | **M (SD)** | **M (SD)** | **M (SD)** |  |  |
| Total Composite Score<sup>b</sup> | 28.34 (12.34) | 41.10 (17.88) | 46.64 (12.01) | F(2,72) = 12.17 | p < .001 |
| Social Cognition | 41.24 (12.70) | 45.95 (14.66) | 50.24 (8.47) | F(2,77) = 3.93 | p = .024 |
| Verbal Learning | 37.12 (7.02) | 42.68 (12.89) | 48.88 (9.02) | F(2,41.92) = 14.53 | p < .001 |
| Visual Learning | 35.06 (17.48) | 38.50 (15.76) | 46.36 (14.34) | H(2) = 6.95 | p = .031 |
| Processing Speed | 36.97 (11.82) | 45.64 (20.37) | 51.76 (13.52) | H(2) = 14.50 | p < .001 |
| Attention<sup>b</sup> | 35.31 (10.20) | 41.05 (13.61) | 46.24 (11.95) | H(2) = 10.95 | p = .004 |
| Working Memory | 36.21 (14.33) | 45.91 (13.19) | 48.84 (9.86) | H(2) = 12.48 | p = .002 |
| Reasoning | 40.94 (9.90) | 47.64 (11.55) | 44.08 (12.30) | F(2,77) = 1.79 | p = .173 |
| **Metabolite Concentration<sup>c</sup>** | **M (SD)** | **M (SD)** | **M (SD)** |  |  |
| NAA | 6.64 (1.36) | 6.13 (1.68) | 6.71 (1.49) | F(2,77) = 1.06 | p = .352 |
| Cr | 7.05 (2.17) | 7.44 (1.10) | 7.52 (1.62) | H(2) = 0.52 | p = .771 |
| Cho | 2.51 (0.70) | 2.48 (0.48) | 2.51 (0.63) | H(2) = 0.10 | p = .952 |
| mI<sup>d</sup> | 4.21 (1.52) | 4.74 (1.36) | 4.85 (1.02) | H(2) = 2.00 | p = .359 |
| Glx<sup>e</sup> | 6.73 (2.97) | 6.86 (2.31) | 7.80 (2.12) | F(2,47) = 0.97 | p = .388 |

Note. <sup>a</sup>For group comparisons, Chi-square test was used for sex; One-way ANOVA for total composite score, social cognition, NAA, and Glx; Welch's ANOVA for verbal learning; and Kruskal-Wallis test for all other variables in the table. <sup>b</sup>Psychosis, n=29; NP-Affective, n=21; healthy controls, n=25. <sup>c</sup>Millimolar. <sup>d</sup>Psychosis, n=21; NP-Affective, n=16; healthy controls, n=20. <sup>e</sup>Psychosis, n=16; NP-Affective, n=16; healthy controls, N=18.

### Primary Group Comparisons in Cognition

ANOVA results are presented in Table 1. There was a significant difference between the psychosis, NP-affective, and healthy control groups in overall cognition (F(2,72)=12.17, p<.001, η^2^_p_=.253). Post-hoc pair-wise comparisons showed that the psychosis group (M=28.34, SD=12.34) had reduced overall cognition performance compared to healthy controls (M=46.64, SD=12.01, p<.001) and NP-affective (M=41.10, SD=17.88, p=.002). Similarly, there was a significant difference between the psychosis, NP-affective, and healthy controls in working memory (H(2)=12.48, p=.002, η^2^_p_=.110), social cognition (F(2,77)=3.93, p=.024, η^2^_p_=.024), visual learning (H(2)=6.95, p=.031, η^2^_p_=.031), verbal learning (F(2,41.92)=14.53, p<.001, η^2^_p_=.219), processing speed (H(2)=14.50, p<.001, η^2^_p_=.136), and attention (H(2)=10.95, p-.004, η^2^_p_=.097), although reasoning scores were similar across groups (F(2,77)=2.40, p=.098, η^2^_p_=.059). Post-hoc pair-wise comparisons showed that psychosis had reduced performance for working memory, social cognition, visual learning, verbal learning, processing speed, and attention compared to healthy controls (p’s≤.010) and working memory compared to NP-affective (p=.016), all unaltered when controlling for age and sex (p’s<.01), except for trend level reductions for social cognition (p=.018) and visual learning (p*=*.020) in the comparison of psychosis and healthy controls.

### Subgroup Analyses for Cognition

There was a significant difference between the schizophrenia, affective psychosis, NP-affective, and healthy controls in overall cognition (F(3,71)=9.64, p<.001, η^2^_p_=.290). Post-hoc pair-wise comparisons showed that schizophrenia (M=24.79, SD=2.86) had reduced overall cognition performance compared to healthy controls (M=46.64, SD=12.01, p<.001) and NP-affective (M=41.10, SD=17.88, p<.001). Similarly, there was a significant difference between the schizophrenia, affective psychosis, NP-affective, and healthy controls in working memory (H(3)=21.65, p<.001, η^2^_p_=.245), social cognition (F(3,76) =4.05, p=.010, η^2^_p_=.138), visual learning (H(3)=9.52, p=.023, η^2^_p_=.086), verbal learning (H(3)=19.17, p<.001, η^2^_p_=.213), processing speed (H(3)=14.88, p=.002, η^2^_p_=.156), and attention (H(3)=11.83, p-.008, η^2^_p_=.181), although reasoning scores were similar across groups (F(3,76)=1.62, p=.190, η^2^_p_=.060). Post-hoc pair-wise comparisons showed that schizophrenia had reduced performance for working memory, social cognition, visual learning, verbal learning, processing speed, and attention compared to healthy controls (p’s<.01), as well as working memory compared to NP-affective and affective psychosis (p’s<.01). Post-hoc comparisons also showed that affective psychosis had reduced performance for verbal learning compared to healthy controls (p=.006).

### Comparisons of Neurometabolites Concentrations Across Groups

The groups did not differ significantly in mean hippocampal metabolite concentrations, both before and after controlling for age and sex (Table 2 shows comparisons between psychosis, NP-affective, and healthy controls). The groups also did not differ significantly in neurometabolite variances.

**Table 2.** Correlations Between Metabolite Biomarker Levels and Cognitive Measures in the Healthy Control, Non-psychosis Affective Disorder, and Psychosis Groups, and Comparisons of Correlations Across Groups.

| **MATRICS** | **Metabolite** | **Psychosis (N=33)<sup>a</sup>** | **NP-Affective (N=22)<sup>a</sup>** | **Healthy Controls (N=25)<sup>a</sup>** | **Psychosis vs. NP-Affective** |  | **Psychosis vs. Healthy Controls** |  | **NP-Affective vs. Healthy Controls** |  |
|---|---|---|---|---|---|---|---|---|---|---|
|  |  | **r<sub>s</sub>** | **r<sub>s</sub>** | **r<sub>s</sub>** | **z** | **p** | **z** | **p** | **z** | **p** |
| **Total Composite Score** | NAA | .014 | .161 | **.414\*** | -0.48 | .628 | -1.47 | .141 | 0.87 | .382 |
|  | Cr | .190 | -.093 | .233 | 0.93 | .352 | -0.16 | .876 | 1.04 | .298 |
|  | Cho | .248 | -.350 | .232 | 2.02 | .043 | 0.06 | .953 | 1.89 | .058 |
|  | mI | .192 | .365 | -.010 | -0.52 | .605 | 0.60 | .546 | -1.07 | .287 |
|  | Glx | .154 | .373 | .143 | -0.56 | .574 | 0.03 | .978 | -0.65 | .513 |
| **Social Cognition** | NAA | -.053 | -.122 | .121 | 0.24 | .812 | -0.62 | .534 | 0.78 | .436 |
|  | Cr | -.117 | -.294 | .296 | 0.63 | .527 | -1.51 | .132 | 1.94 | .052 |
|  | Cho | .116 | -.152 | .051 | 0.92 | .358 | 0.23 | .816 | 0.65 | .514 |
|  | mI | .149 | .049 | .163 | 0.28 | .781 | -0.04 | .966 | 0.31 | .754 |
|  | Glx | .125 | .364 | -.038 | -0.65 | .514 | 0.43 | .666 | -1.11 | .268 |
| **Verbal Learning** | NAA | .110 | .051 | .269 | 0.20 | .839 | -0.59 | .556 | 0.72 | .473 |
|  | Cr | .129 | -.083 | .121 | 0.73 | .468 | 0.03 | .977 | 0.65 | .513 |
|  | Cho | .191 | **-.446\*** | .219 | 2.30 | .021 | -0.10 | .917 | 2.24 | .025 |
|  | mI | .312 | **.503\*** | .069 | -0.63 | .526 | 0.75 | .453 | -1.31 | .189 |
|  | Glx | .227 | .387 | .114 | -0.45 | .651 | 0.31 | .758 | -0.78 | .438 |
| **Visual Learning** | NAA | .127 | -.071 | -.114 | 0.68 | .498 | 0.86 | .388 | -0.14 | .890 |
|  | Cr | .057 | -.388 | -.178 | 1.59 | .112 | 0.84 | .399 | 0.73 | .464 |
|  | Cho | **.514\*** | -.405 | -.178 | 3.40 | <.001 | 2.67 | .008 | 0.80 | .425 |
|  | mI | .118 | -.008 | -.362 | 0.35 | .728 | 1.47 | .141 | -1.01 | .314 |
|  | Glx | .140 | .118 | -.046 | 0.06 | .955 | 0.49 | .622 | -0.43 | .664 |
| **Processing Speed** | NAA | .114 | .104 | .168 | 0.03 | .972 | -0.20 | .844 | 0.21 | .835 |
|  | Cr | .014 | .044 | -.051 | -0.10 | .918 | 0.23 | .817 | -0.30 | .761 |
|  | Cho | .151 | -.258 | -.065 | 1.42 | .156 | 0.77 | .439 | 0.63 | .525 |
|  | mI | .279 | .458 | -.001 | -0.57 | .567 | 0.85 | .395 | -1.35 | .178 |
|  | Glx | .090 | .449 | .053 | -1.00 | .316 | 0.10 | .922 | -1.14 | .256 |
| **Attention** | NAA | -.041 | .295 | .310 | -1.13 | .260 | -1.25 | .212 | 0.05 | .959 |
|  | Cr | -.026 | .113 | .120 | -0.45 | .649 | -0.51 | .613 | 0.02 | .982 |
|  | Cho | -.016 | -.200 | .122 | 0.61 | .543 | -0.48 | .632 | 1.02 | .306 |
|  | mI | -.184 | .374 | -.328 | -1.59 | .112 | 0.46 | .648 | -1.99 | .047 |
|  | Glx | -.264 | .346 | .176 | -1.50 | .133 | -1.10 | .272 | -0.48 | .629 |
| **Working Memory** | NAA | -.165 | .086 | **.422\*** | -0.86 | .389 | - 2.20 | .028 | 1.16 | .245 |
|  | Cr | -.179 | -.021 | .212 | -0.55 | .585 | -1.41 | .158 | 0.75 | .451 |
|  | Cho | .071 | -.369 | .254 | 1.56 | .118 | -0.67 | .502 | 2.09 | .037 |
|  | mI | .176 | .256 | .242 | -0.23 | .818 | -0.20 | .838 | -0.04 | .968 |
|  | Glx | .041 | .164 | .045 | -0.32 | .751 | -0.01 | .992 | -0.32 | .751 |
| **Reasoning** | NAA | .131 | -.030 | **.433\*** | 0.55 | .581 | -1.18 | .237 | 1.58 | .115 |
|  | Cr | -.086 | -.147 | .340 | 0.21 | .833 | -1.57 | .117 | 1.60 | .109 |
|  | Cho | .325 | -.360 | .382 | 2.44 | .015 | -0.23 | .816 | 2.49 | .013 |
|  | mI | .073 | .196 | .230 | -0.34 | .730 | -0.48 | .634 | 0.10 | .923 |
|  | Glx | -.094 | .213 | .205 | -0.79 | .428 | -0.80 | .425 | -0.02 | .982 |

Note. <sup>\*</sup>Significant within-group correlations (α=0.05) are shown in bold and marked with an asterisk. <sup>a</sup>Some analyses had fewer participants because of missing values for mI and Glx and/or attention and overall cognition, as detailed in Supplementary Material. r<sub>s</sub> = Spearman’s correlation coefficient.

### Neurometabolite Concentrations and Cognition by Groups

Although neurometabolite levels were similar across groups, we found both significant with-group correlations between neurometabolite levels and cognitive domains and significant between-group differences in correlation coefficients (Tables 2 and 3).

**Table 3.** Correlations Between Metabolite Biomarker Levels and Cognitive Measures in the Schizophrenia and Affective Psychosis Groups, and Comparisons of Correlations Across Groups.

|  |  | Schizophrenia (N=20)<sup>a</sup> | Affective Psychosis (N=13)<sup>a</sup> | Schizophrenia vs Affective Psychosis |  | Schizophrenia vs. NP-Affective<sup>b</sup> |  | Schizophrenia vs. Healthy Controls<sup>c</sup> |  | Affective Psychosis vs. NP-Affective<sup>b</sup> |  | Affective Psychosis vs. Healthy Controls<sup>c</sup> |  |
|---|---|---|---|---|---|---|---|---|---|---|---|---|---|
|  |  | r<sub>s</sub> | r<sub>s</sub> | z | p | z | p | z | p | z | p | z | p |
| **Total Compositive Score** | NAA | -.030 | -.030 | 0.00 | 1.000 | 0.56 | .575 | 1.43 | .152 | 0.43 | .666 | 1.08 | .278 |
|  | Cr | .387 | -.176 | 1.29 | .196 | -1.46 | .144 | -0.52 | .603 | 0.19 | .849 | 0.96 | .339 |
|  | Cho | .267 | -.193 | 1.04 | .301 | -1.86 | .063 | -0.11 | .910 | -0.38 | .703 | 0.99 | .320 |
|  | mI | .275 | -.036 | 0.55 | .586 | 0.25 | .806 | -0.76 | .450 | 0.73 | .464 | 0.05 | .963 |
|  | Glx | .183 | .800 | -0.85 | .398 | 0.42 | .675 | -0.09 | .932 | -0.68 | .496 | -0.92 | .355 |
| **Social Cognition** | NAA | .178 | **-.581\*** | 2.12 | .034 | -0.91 | .365 | -0.18 | .857 | 1.39 | .166 | 2.06 | .039 |
|  | Cr | .250 | -.457 | 1.88 | .060 | -1.67 | .094 | 0.15 | .878 | 0.49 | .626 | 2.09 | .037 |
|  | Cho | .313 | **-.575\*** | 2.46 | .014 | -1.43 | .153 | -0.84 | .398 | 1.28 | .199 | 1.85 | .064 |
|  | mI | .480 | -.739 | 2.52 | .012 | -1.16 | .247 | -0.93 | .354 | 1.74 | .081 | 2.00 | .046 |
|  | Glx | **.673\*** | -.580 | 2.14 | .032 | -0.93 | .354 | -1.87 | .062 | 1.63 | .103 | 0.99 | .323 |
| **Verbal Learning** | NAA | .203 | -.073 | 0.70 | .484 | -0.46 | .643 | 0.22 | .829 | 0.32 | .751 | 0.91 | .360 |
|  | Cr | .304 | -.032 | 0.87 | .385 | -1.19 | .234 | -0.60 | .551 | -0.13 | .896 | 0.40 | .687 |
|  | Cho | .222 | .025 | 0.50 | .614 | -2.11 | .035 | -0.01 | .992 | -1.29 | .196 | 0.52 | .604 |
|  | mI | .296 | .429 | -0.26 | .793 | 0.61 | .545 | -0.61 | .542 | 0.17 | .869 | -0.70 | .483 |
|  | Glx | .274 | .200 | 0.11 | .909 | 0.27 | .786 | -0.36 | .716 | 0.32 | .748 | -0.14 | .889 |
| **Visual Learning** | NAA | .193 | -.119 | 0.79 | .429 | -0.80 | .425 | -0.96 | .337 | 0.12 | .901 | 0.01 | .989 |
|  | Cr | .403 | -.416 | 2.18 | .029 | -2.51 | .012 | -1.88 | .060 | 0.09 | .932 | 0.69 | .491 |
|  | Cho | **.543\*** | .368 | 0.56 | .577 | -3.11 | .002 | -2.44 | .015 | -2.09 | .037 | -1.48 | .138 |
|  | mI | .322 | -.036 | 0.63 | .526 | -0.83 | .404 | -1.84 | .065 | 0.05 | .961 | -0.62 | .537 |
|  | Glx | .358 | .058 | 0.46 | .646 | -0.55 | .585 | -0.92 | .358 | 0.09 | .925 | -0.16 | .869 |
| **Processing Speed** | NAA | .039 | .153 | -0.29 | .773 | 0.20 | .845 | 0.40 | .686 | -0.13 | .899 | 0.04 | .968 |
|  | Cr | .062 | -.026 | 0.22 | .825 | -0.05 | .957 | -0.35 | .726 | 0.18 | .858 | -0.07 | .948 |
|  | Cho | .006 | .354 | -0.91 | .361 | -0.81 | .419 | -0.22 | .826 | -1.62 | .105 | -1.14 | .254 |
|  | mI | .139 | .519 | -0.75 | .456 | 0.87 | .386 | -0.36 | .716 | -0.14 | .888 | -1.04 | .300 |
|  | Glx | .301 | -.029 | 0.49 | .623 | 0.37 | .712 | -0.56 | .574 | 0.80 | .424 | 0.13 | .897 |
| **Attention** | NAA | -.069 | -.104 | 0.08 | .938 | 1.09 | .277 | 1.19 | .236 | 0.92 | .359 | 0.98 | .327 |
|  | Cr | .121 | -.352 | 1.08 | .280 | -0.02 | .981 | -0.00 | .998 | 1.08 | .280 | 1.13 | .260 |
|  | Cho | .033 | -.244 | 0.62 | .534 | -0.69 | .493 | 0.27 | .785 | 0.10 | .917 | 0.86 | .392 |
|  | mI | .121 | **-.847\*** | 2.34 | .019 | 0.66 | .508 | -1.19 | .232 | 2.87 | .004 | 1.63 | .103 |
|  | Glx | -.017 | -.800 | 1.00 | .317 | 0.77 | .444 | 0.40 | .687 | 1.41 | .160 | 1.24 | .216 |
| **Working Memory** | NAA | -.206 | -.117 | -0.23 | .818 | 0.88 | .377 | 2.04 | .041 | 0.52 | .602 | 1.49 | .137 |
|  | Cr | .022 | -.037 | 0.15 | .882 | -0.13 | .897 | 0.60 | .550 | 0.04 | .967 | 0.66 | .508 |
|  | Cho | -.036 | .029 | -0.16 | .870 | -1.05 | .293 | 0.92 | .360 | -1.07 | .287 | 0.60 | .545 |
|  | mI | .030 | .571 | -1.06 | .289 | 0.57 | .571 | 0.56 | .575 | -0.68 | .498 | -0.72 | .469 |
|  | Glx | .322 | -.086 | 0.61 | .543 | -0.36 | .719 | -0.63 | .528 | 0.39 | .694 | 0.21 | .836 |
| **Reasoning** | NAA | -.056 | .444 | -1.34 | .181 | 0.08 | .938 | 1.61 | .108 | -1.30 | .194 | -0.04 | .972 |
|  | Cr | -.143 | -.047 | -0.24 | .808 | -0.01 | .990 | 1.54 | .123 | -0.26 | .796 | 1.05 | .293 |
|  | Cho | .069 | **.729\*** | -2.15 | .032 | -1.34 | .182 | 1.03 | .302 | -3.34 | <.001 | -1.37 | .169 |
|  | mI | -.101 | .393 | -0.88 | .376 | 0.73 | .464 | 0.87 | .386 | -0.38 | .705 | -0.33 | .744 |
|  | Glx | -.285 | .429 | -1.09 | .276 | 1.09 | .277 | 1.09 | .274 | -0.38 | .705 | -0.40 | .692 |

Note. <sup>\*</sup>Significant within-group correlations (α=0.05) are shown in bold and marked with an asterisk. <sup>a</sup>Some analyses had fewer participants because of missing values for mI and Glx and/or attention and overall cognition, as detailed in Supplementary Materials. <sup>b</sup>See Table 2 for correlation coefficients for the NP-Afective group. <sup>c</sup>See Table 2 for correlation coefficients for the healthy control group. r<sub>s</sub> = Spearman's correlation coefficient.

### Healthy controls (HC)

demonstrated lower concentrations of the neuronal integrity biomarker (NAA) in association with lower scores for overall cognition (r_s_=.414, *p=*.039), reasoning (r_s_=.433, *p=*.031) and working memory (r_s_=.422, *p=*.036), and the association with working memory in healthy controls was opposite in direction from the association between the same measures in psychosis and schizophrenia (*p’s*<.05).

### Non-psychotic affective group (NP-affective)

showed a significant association of lower verbal learning scores with reduced levels of the glia biomarker (mI; r_s_=.503, p=.047) and increased levels of the membrane/myelin turnover biomarker (r_s_=−.446, *p=*.038). The association with membrane/myelin turnover was opposite from healthy controls, psychosis, and schizophrenia (*p’s*<.05).

### Psychotic group (Psy)

demonstrated a significant association between reduced levels of the membrane/myelin turnover biomarker (Cho) and visual learning deficits (r_s_=.514, *p=*.002), which was opposite from NP-affective (p<.001) and healthy controls (p=.008).

### Separated schizophrenia and affective psychosis (affective psychosis) subgroups

Different profiles of metabolite abnormalities were shown for social cognition. Social cognition impairments were related to reduced levels of the marker for glutamatergic neurotransmission (Glx) in schizophrenia (r_s_=.673, p=.033), but to elevated concentrations of the markers for membrane/myelin turnover (r_s_=−.575, p=.040) and neuronal integrity (r_s_=−.581, p=.037) in affective psychosis. Moreover, the associations of social cognition with all of the metabolite biomarkers except for Cr were opposite across the two groups (*p’s*<.05).

Additional significant findings across the psychosis subgroups included that affective psychosis showed increased levels of the gliosis marker in association with attention deficits (r_s_=−.847, p=.016) and reduced levels of the membrane/myelin turnover marker in association with lower reasoning scores (r_s_=.729 p=.005), both of which were opposite in schizophrenia and NP-affective (*p’s*<.05). Also, schizophrenia showed an association between decreased levels of the membrane/myelin turnover marker and visual learning impairments (r_s_=.543, p=.013), similar to the combined psychosis group. The association in schizophrenia was opposite from NP-affective (p=.002) and healthy controls (p=.015).

### Effects of Age and Illness Duration on Neurometabolite-Cognition Associations

We found several significant correlations of age and illness duration with metabolite levels and cognition in the psychotic groups, but none of these explained any of the associations between cognitive scores and metabolite levels in these groups (see Supplementary Results). As expected, illness duration was strongly associated with age in psychosis (r_s_=.861, p<.001), schizophrenia (r_s_=.824, *p*<. 001), and affective psychosis (r_s_=.875, *p*<.001).

## DISCUSSION

This study examined associations between cognitive measures and hippocampal metabolite concentrations in healthy participants, those with non-psychosis affective disorders, and psychotic participants to investigate the cellular and molecular underpinnings of cognitive deficits. Cognitive scores were similar for healthy controls and non-psychotic affective disorders, with both performing significantly better than the psychotic group for overall cognition. Healthy controls also consistently outperformed the psychotic group across the MATRICS cognitive domains. While mean metabolite concentrations were similar across groups, the within-group mapping of metabolites to cognitive domains suggested distinct associations between cellular pathology and cognitive deficits across diagnostic classifications.

Lower overall cognition, working memory, and reasoning scores were associated with reduced levels of the neuronal integrity biomarker (NAA) in healthy controls, none of which was observed in the clinical groups. Because NAA is primarily synthesized in neuronal mitochondria, decreased levels could reflect changes in neuronal metabolic function without necessarily indicating neuronal loss or degeneration. ^25^ This possibility may be particularly relevant in otherwise healthy individuals, in whom such pathology would be less expected. Alternatively, if reflective of reduced neuronal integrity, lower NAA levels in the controls could reflect the effects of adverse life-course exposures^26^ or genetic variations. Aging and neurodegenerative disorders also compromise neuronal integrity, ^27^ although these are less likely explanations given that our study groups were not aged. Additionally, systemic inflammation, metabolic disorders, obesity, and cardiovascular risk factors may be relevant to reduced neuronal integrity. ^28^ These conditions are associated with physiological processes that can adversely impact brain health.

Among non-psychotic affective cases, reduced verbal memory scores were associated with elevated levels of the biomarkers for membrane/myelin turnover (Cho) and glia (mI), and the association with Cho was opposite in direction from that observed in healthy and psychotic participants. A profile of elevated choline and reduced myo-inositol could indicate a process characterized by increased membrane turnover (such as inflammation or demyelination) with concurrent glial dysfunction or loss. Neuroinflammatory exposures are commonly described in mood disorders, ^29, 30^ and some findings implicate demyelinating processes in these illnesses. ^31^ Early-life stress may be particularly relevant to neuroinflammatory processes and their effects on hippocampal cellular integrity in mood disorders. These patients report increased rates of childhood trauma, which has been linked to alterations in their stress-response and inflammatory systems. ^29^ Chronic activation of the hypothalamic-pituitary-adrenal (HPA) axis and excessive glucocorticoid exposure induced by early-life stress can cause hippocampal neurotoxicity. ^32^ Stress can also activate and dysregulate glial cells, including microglia and astrocytes, ^33^ potentially promoting inflammatory processes and elevated membrane turnover, the latter reflected in elevated MRSI signals from choline-containing compounds, such as glycerophosphocholine and phosphocholine. ^8, 34^ Also of relevance to our findings, increased membrane/myelin turnover signals in depression have been linked to a high past illness burden, ^34^ as is typical of the depression cases recruited from our hospital settings.

In contrast to non-psychotic affective cases, in the combined schizophrenia and affective psychosis group reduced (rather than increased) membrane/myelin turnover was related to cognitive deficits, specifically to visual learning deficits. The same association was also observed in schizophrenia alone, and both associations were opposite in direction to those observed in the non-psychotic groups. The finding in schizophrenia may suggest a hypomyelination process arising from abnormalities in oligodendrocyte function, consistent with schizophrenia studies showing white matter deficits and reduced expression of genes and proteins associated with myelin and oligodendrocytes. ^35^ Myelin is essential for efficient neural signal conduction, and its reduction may therefore contribute to cognitive deficits.

Separating the psychosis group into schizophrenia and affective psychosis revealed strikingly distinct neurometabolite biomarker profiles associated with social cognition, reflected in both the metabolites showing significant associations and the broader pattern of between-group differences. Social cognition deficits were associated with biomarkers of reduced glutamatergic neurotransmission (Glx) in schizophrenia and increased membrane/myelin turnover (Cho) and neural integrity (NAA) in affective psychosis, and associations with social cognition were opposite in direction and significantly different between the groups for every quantitated metabolite biomarker except Cr. This pattern of results suggests a diagnosis-dependent double dissociation of the neurochemical substrate of social cognition in schizophrenia versus affective psychosis rather than a single shared substrate across psychotic disorders. These findings are particularly noteworthy because social cognition is more predictive of functional outcome than any other cognitive ability among primary psychoses. ^36^ In light of this, our results suggest distinct pathways to poor outcomes for patients with schizophrenia and affective psychosis who have social cognition deficits.

The association between reduced Glx and social cognition deficits in schizophrenia may reflect disturbances in excitatory transmission of the kind proposed to contribute to schizophrenia symptoms through NMDA receptor hypofunction. ^37^ Dysregulated glutamatergic neurotransmission in the hippocampus and other brain regions may interfere with the coordination of neural circuits supporting higher-order cognitive functions. ^38^ The associations of decreased Glx and Cho with different cognitive domains in schizophrenia may together reflect broader, interconnected disturbances in neuronal and cellular function in the illness. Oxidative stress may contribute to these abnormalities through disruption of glutamatergic signaling and alterations in phospholipid homeostasis. ^39, 40^ In contrast to schizophrenia, social cognition deficits in affective psychosis were related to increased Cho and NAA, potentially reflecting increased membrane turnover alongside altered neuronal metabolism. Neuroinflammation may contribute to the increased membrane turnover, as also proposed above for the elevated Cho associated with verbal learning deficits in non-psychotic affective cases. This interpretation is consistent with numerous findings showing elevated inflammatory cytokines in both psychotic disorders and affective disorders, ^41–43^ although few studies have examined affective psychoses separately. One of the few found increased levels of inflammatory cytokines (including IL-1β, IL-6, IFNγ, IL-17, and IL-15) in psychotic affective disorders compared to healthy controls across both baseline and immune activation conditions. ^43^ More compelling evidence suggestive of neuroinflammation in our affective psychosis group comes from the strong association between attention deficits and increased levels of mI, a glial biomarker closely linked to astrocytic function and inflammatory processes. ^8^ As such, neuroinflammatory mechanisms may contribute to deficits across multiple cognitive domains in affective psychoses. The counterintuitive association of social cognition deficits with increased NAA in our study could reflect a compensatory change in neuronal metabolism or energetic capacity in response to underlying neural dysfunction. A previous study similarly linked elevated NAA to reduced cognitive performance in healthy older adults and interpreted this finding as potentially reflecting increased mitochondrial energy capacity. ^44^

A different pattern of Cho associations in affective psychosis emerged for reasoning/problem solving, with lower scores associated with reduced membrane/myelin turnover. As proposed above for schizophrenia, reduced membrane/myelin turnover in affective psychosis may suggest a process in which oligodendrocyte dysfunction leads to white matter deficits through decreased myelin production. Thus, different cognitive deficits in affective psychosis were associated with opposing patterns of Cho abnormalities, potentially reflecting distinct underlying pathologies.

This is the first hippocampal MRS study to measure and compare associations between metabolite levels and cognition across different psychiatric classifications, including different psychotic disorders. This approach revealed distinct associations in schizophrenia and affective psychosis that were largely obscured when all psychotic cases were intermixed, particularly for social cognition, suggesting that DSM-5 classification of these disorders remains meaningful despite transdiagnostic conceptualizations of psychosis. Additional strengths of our study include the systematic ascertainment of stable consecutive psychiatric cases from the same hospital system, rigorously evaluated to minimize confounding by medication, ascertainment approaches, diagnostic practices. The analysis also accounted for age and disease duration, which did not explain any of the associations in the psychotic groups. Additionally, previous MRS studies of psychosis commonly used metabolite ratios rather than absolute hippocampal concentrations, which may help to explain the highly variable results across the studies. Finally, our novel approach to quantifying metabolites using whole-hippocampal multi-voxel MRSI was developed to overcome methodological weakness in the field, with support from an earlier NIMH “challenge grant” awarded to DM and OG. Most prior MRS studies used single-voxel spectroscopy, which cannot account for the hippocampus’ irregular shape or CSF/gray/white matter partial volume effects and is usually unilateral, whereas our multi-voxel MRS covered the entire 3-dimensional hippocampus with precision.

Our study also has notable weaknesses. While we recruited and comprehensively evaluated the 80 planned participants, the individual diagnostic groups were more modest number, particularly after the psychosis cohort was divided into schizophrenia and affective psychosis. Also, this was not longitudinal research, although it was part of a multi-level cross sectional analysis that included rich symptom assessments, autonomic function, and gut and oral microbiota. Next, similar to other studies that employ MRS localization, we focused only on bilateral hippocampus, and as such our analysis did not account for hippocampal laterality. Furthermore, while all of our subjects were on stable medications and in active treatment, some had long, 20-plus-year disease durations, and so accounting for all medications and subject compliance in the elapsed decades was not possible. A final but important weakness is that our samples were too small for sex stratification. Although sex distributions did not differ significantly across groups, there were still noticeable differences in male to female ratio for the combined psychosis and schizophrenia groups compared to the healthy control and non-psychotic affective disorder groups. We therefore cannot completely rule out the possibility that sex influenced the findings of our correlation analyses. However, the differences in sex distribution across groups reflect the proportions observed in treatment through our systematic ascertainment.

This analysis of MRSI neurometabolite biomarkers and cognitive performance suggests distinct cellular pathologies underlying cognitive deficits across psychotic disorders, non-psychotic affective disorders, and healthy persons. Most notably, different neurometabolite biomarkers were associated with social cognition deficits in schizophrenia versus affective psychosis. These findings challenge transdiagnostic approaches such as the NIMH RDoC framework, which posit similar cellular mechanisms for deficits in a given cognitive domain that cut across diagnostic categories and even manifest at subclinical levels in the healthy population. It may therefore be premature to dismiss DSM categories and similar conceptualizations in the ICD or to conclude that all psychoses belong in a single classification. Ultimately, our findings highlight the potential for interventions tailored to different psychotic disorders, particularly those targeting social cognition, which has strong predictive power for real-world functional outcomes.

## ACKNOWLEDGEMENTS

This work was supported by NIMH 1R01MH110418-01A1 (DM), NIBIB P41 EB017183 (OG), and NIBIB U24 EB02898 (OG).

