## Supplementary Information for "Distinct Hippocampal Cellular Pathologies Influence Cognition Across Diagnostic Categories and Distinguish Schizophrenia from Affective Psychosis"

**SUPPLEMENTARY RESULTS**

Some participants had missing values for mI and Glx because the signal for these neurometabolites did not meeting quality control criteria outlined in the text. The peaks for mI and Glx in the MRS spectrum are much smaller than the peaks for the other metabolites measured; consequently, the signal-to-noise ratios (SNRs) for mI and Glx sometimes fall below acceptable thresholds. Additionally, several participants had missing MATRICS attention and overall cognition scores because they did not complete the attention test. Specifically, in the psychosis group, 29 participants had attention and overall cognition values, including 19 with schizophrenia and 10 with affective psychosis; 16 had Glx values, including 10 with schizophrenia and 6 with affective psychosis; and 21 had mI values, including 14 with schizophrenia and 7 with affective psychosis. Furthermore, for correlation analyses examining relationships between Glx and either attention or overall cognition in the psychosis group, data were available for 13 participants, including 9 with schizophrenia and 4 with affective psychosis. In the NP-Affective group, 21 participants had attention and overall cognition scores, 16 had Glx values, and 16 had mI values. In the healthy control group, 18 participants had Glx values and 20 had mI values.

Covariates for age and illness duration in the psychotic groups. In the combined group of schizophrenia and affective psychosis, older age was related to poorer working memory (r_s_=−.351, p=.045), a cognitive domain not related to any metabolites in this group. In affective psychosis, older age was associated with increased levels of the membrane/myelin turnover biomarker (Cho) (r_s_=.670, p=.012). However, the correlations of social cognition with age and illness duration in affective psychosis were weak and not significant, and overall Cho levels in this group were similar to that in the other groups both before and after controlling for age and sex. This suggests that the effects of older age did not explain the association between increased Cho and poorer social cognition in affective psychosis. Also in affective psychosis, longer illness duration was associated worse overall cognition (r_s_=−.744, p=.014), but neither illness duration nor age were significantly related to any of the seven cognitive domains that contribute to the overall cognition score. The association between illness duration and overall cognition was mostly explained by non-significant moderate associations with reduced verbal memory and working memory, neither of which were associated with metabolites in affective psychosis. No significant associations of age or illness duration with cognition or metabolite levels were found in schizophrenia.

**SUPPLEMENTARY TABLES**

**Supplementary Table 1. Demographic, Cognitive, and Imaging Data for the Psychosis Group, Stratified by Diagnosis**

|  | **Schizophrenia  (n = 20)** | **Affective  Psychosis  (n = 13)** | **Test Statistic**^ab^ | **p** |
| --- | --- | --- | --- | --- |
| **Demographics** |  |  |  |  |
| Age, *M (SD)* | 38.20 (2.40) | 32.08 (2.92) | U = 84.50 | p = .093 |
| Disease duration (years), *M (SD)* | 20.05 (2.83) | 16.08 (3.51) | t(31) = 0.88 | p = .385 |
| Sex *(Percent Male:Female)* | 60.0 : 40.0 | 38.5 : 61.5 | χ²(1) = 1.46 | p = .226 |
| **MATRICS Cognitive Battery** | **M (SD)** | **M (SD)** |  |  |
| Total Composite Score^c^ | 24.79 (2.86) | 35.10 (2.92) | t(27) = -2.30 | p = .030 |
| Social Cognition | 37.90 (2.79) | 46.38 (3.25) | t(31) = -1.96 | p = .060 |
| Verbal Learning | 35.95 (1.62) | 38.92 (1.81) | t(31) = -1.20 | p = .241 |
| Visual Learning | 31.15 (3.99) | 41.08 (4.36) | U = 175.00 | p = .097 |
| Processing Speed | 35.85 (2.50) | 38.69 (3.60) | U = 155.00 | p = .356 |
| Attention^b^ | 33.58 (2.50) | 38.60 (2.63) | t(27) = -1.27 | p = .213 |
| Working Memory | 29.85 (2.93) | 46.00 (2.84) | t(31) = -3.76 | p < .001 |
| Reasoning | 40.40 (2.17) | 41.77 (2.92) | t(31) = 0.38 | p = .704 |
| **Metabolite Concentration^d^** | **M (SD)** | **M (SD)** |  |  |
| NAA | 6.69 (.332) | 6.57 (.331) | U = 120.50 | p = .726 |
| Cr | 7.56 (.474) | 6.26 (.581) | t(31) = 1.73 | p = .094 |
| Cho | 2.44 (.161) | 2.62 (.186) | U = 156.50 | p = .328 |
| mI^e^ | 4.02 (.386) | 4.60 (.643) | U = 54.00 | p = .709 |
| Glx^f^ | 7.54 (1.01) | 5.39 (.888) | t(14) = 1.45 | p = .168 |

^a^Tests in this table only compared schizophrenia and affective psychosis, whereas in the main text these groups were also compared to healthy controls and non-psychotic affective disorders using ANOVA. ^b^For group comparisons in the table, Mann-Whitney U test was used for age, visual learning, verbal learning, NAA, Cho, and mI; Chi-square test for sex; and T-test for all other variables in the table. ^c^Schizophrenia, n=19; Affective Psychosis, n=10. ^d^Millimolar. ^e^Schizophrenia, n=14; Affective Psychosis, n=7. ^f^Schizophrenia, n=10; Affective Psychosis, n=6.
